# Cryo-EM structures of the ATP release channel pannexin 1

**DOI:** 10.1101/2020.01.05.895235

**Authors:** Zengqin Deng, Zhihui He, Grigory Maksaev, Ryan M. Bitter, Michael Rau, James A.J. Fitzpatrick, Peng Yuan

**Author notes:** These authors contributed equally to this work (Zengqin Deng, Zhihui He). Correspondence should be addressed to P.Y.

## Abstract

The plasma membrane ATP release channel pannexin 1 has been implicated in numerous physiological and pathophysiological processes associated with purinergic signaling, including cancer progression, apoptotic cell clearance, inflammation, blood pressure regulation, oocyte development, epilepsy and neuropathic pain. Here, we present near-atomic resolution structures of *Xenopus tropicalis* and *Homo sapiens* PANX1 determined by cryo-electron microscopy that reveal a heptameric channel architecture. Compatible with ATP permeation, the transmembrane pore and cytoplasmic vestibule are exceptionally wide. An extracellular tryptophan ring located at the outer pore creates a constriction site, potentially functioning as a molecular sieve that restricts the size of permeable substrates. In combination with functional characterization, this work elucidates the previously unknown architecture of pannexin channels and establishes a foundation for understanding their unique channel properties as well as for developing rational therapies.

## Introduction

Adenosine triphosphate (ATP), in addition to its classical role as the energy currency of living cells, is an important extracellular ligand for numerous cellular signaling events. Broadly expressed, the plasma membrane ATP release channel pannexin 1 (PANX1) has been implicated in a multitude of biological processes associated with purinergic signaling^1,2^. For instance, PANX1 mediates nucleotide release from apoptotic cells during apoptosis^3,4^, regulates blood pressure in vascular smooth muscle^5,6^, promotes cancer progression^7,8^, and contributes to mechanical allodynia^9^, chronic epilepsy^10^, neurological disorders and pain^11–14^. More recently, genetic mutations in PANX1 have been reported to cause abnormal oocyte development leading to female infertility^15^. The expanding list of physiological and pathophysiological functions positions PANX1 as a promising therapeutic target for the treatment of numerous disease conditions, including cancer, epilepsy, infertility and neuropathic pain.

In accordance with a broad spectrum of biological functions, PANX1 is reportedly activated by remarkably diverse physiological mechanisms^1,2,16^, including signaling pathways mediated by membrane receptors^6,11,17–21^, elevated extracellular potassium^22–24^ or intracellular calcium levels^17^, proteolytic cleavage by caspases^3,25,26^, hypoxemia^27^, and membrane stretch and voltage^8,24,28–31^. More strikingly, previous studies suggest that different modes of activation result in distinct open conformations with varying permeability and unitary conductance^1,2,23,26^. Certain physiological stimuli have been reported to induce an expanded open conformation with a high unitary conductance of ∼500 pS^21,28,29,32^, permeable to standard ions such as Na^+^, K^+^, Cl^-^ as well as large molecules such as ATP and fluorescent dyes^28,29,32^. However, caspase and voltage activations render a markedly reduced single-channel conductance (<100 pS)^26,31^. Additionally, sequential removal of the distal C-terminal tail of individual subunits in the oligomeric channel gives rise to gradual increase in unitary conductance, open probability and molecular dimension of permeants^26^. Understanding these perplexing and even contradicting channel properties represents a remarkable challenge without high-resolution structural information of the channel protein. To overcome this obstacle, here we describe near-atomic resolution structures of *Xenopus tropicalis* and *Homo sapiens* PANX1 determined by single-particle cryo-electron microscopy (cryo-EM). Together with functional characterization, our results provide insights into substrate permeability and channel gating as well as a template for further mechanistic investigation and structure-informed rational therapies targeting PANX1 for the treatment of numerous pathological conditions.

## Results

### Structure determination of a full-length functional PANX1 channel

We evaluated multiple PANX1 orthologs and on the basis of biochemical stability decided to focus our structural studies on the *Xenopus tropicalis* channel (xPANX1), which shares 65% sequence identity with the human protein (Supplementary Fig. 1). In excised inside-out membrane patches, xPANX1 channels are spontaneously activated minutes following patch excision, and the currents decrease with application of the PANX1 inhibitor carbenoxolone (CBX) to the bath solutions, i.e., the intracellular side of the plasma membrane (Fig. 1a-d). Current reduction is more pronounced at negative membrane potential (Supplementary Fig. 2a, b). In symmetrical 150 mM NaCl and 150 mM KCl conditions, xPANX1 channels exhibit similar properties, including current-voltage relationship, CBX inhibition, and single-channel conductance (23.0 ± 3.8 pS in NaCl and 24.7 ± 5.3 pS in KCl at -60 mV membrane potential) (Fig. 1a-d, Supplementary Fig. 2a, b). Thus, in excised membrane patches, xPANX1 displays a small unitary conductance and is not further activated by extracellular K^+^.

**Figure 1.**
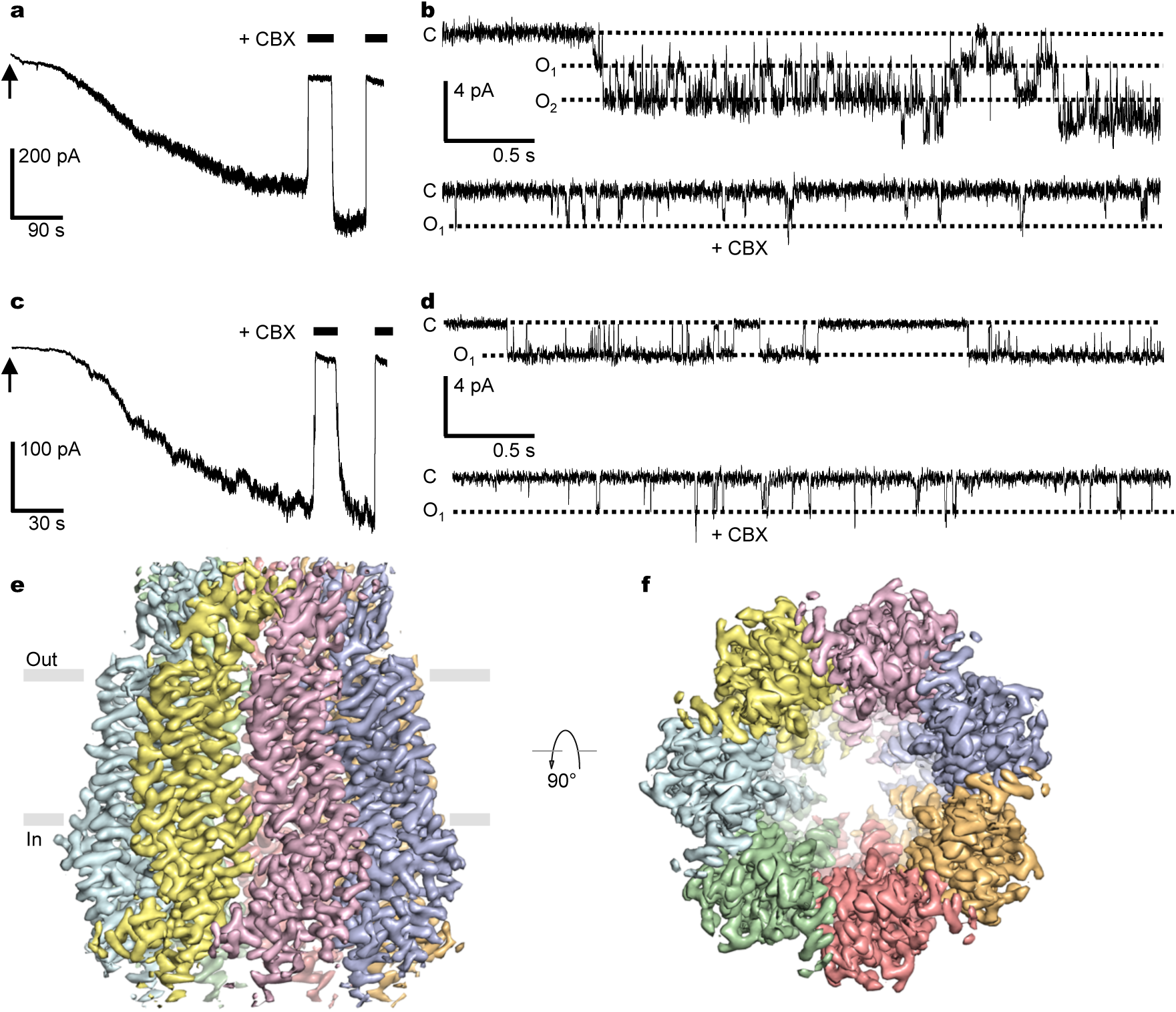
Electrophysiology and cryo-EM reconstruction of xPANX1. **a**, xPANX1-mediated current in an excised membrane patch recorded in symmetrical 150 mM NaCl at -30 mV. Application of the channel inhibitor carbenoxolone (CBX, 100 µM) is indicated by black bars. The arrow indicates patch excision. **b**, Unitary currents of xPANX1 (single-channel conductance 23.0 ± 3.8 pS (mean ± s.d.), 5 independent biological experiments) in 150 mM NaCl at -60 mV in the absence (top panel) and presence (bottom panel) of 100 µM CBX. **c**, xPANX1-mediated current of xPANX1 in an excised membrane patch recorded in symmetrical 150 mM KCl at -30 mV. **d**, Unitary currents of xPANX1 (single-channel conductance 24.7 ± 5.3 pS (mean ± s.d.), 5 independent biological experiments) in 150 mM KCl at -60 mV in the absence (top panel) and presence (bottom panel) of 100 µM CBX. In all experiments in **a**-**d**, CBX was added to the bath solutions (i.e., from the cytoplasmic side). **e, f**, Cryo-EM density of the full-length xPANX1 channel at 3.4 Å resolution viewed parallel to the membrane (**e**) and from the intracellular side (**f**). The electron density is contoured at 6.0 σ and individual subunits are uniquely colored. Gray lines indicate the membrane boundary.

The full-length xPANX1 channel consisting of 428 amino acids was expressed in *Pichia pastoris*, purified to homogeneity, and subjected to single-particle cryo-EM analysis. Initial 2D classification indicated a heptameric channel assembly, and 3D reconstruction yielded a structure at an overall resolution of ∼3.4 Å with C7 symmetry imposed (Fig. 1e, f, Supplementary Fig. 3, Table 1). The electron density map is of excellent quality and displays side-chain densities for the majority of amino acids (Fig. 1e, f, Supplementary Fig. 4), allowing confident model building, which is guided by bulky side chains and ascertained by disulfide bonds present in the extracellular domain. The refined atomic model fits well into the electron density with good geometry (Supplementary Fig. 4 and Table 1). The amino- and carboxyl-terminal portions (amino acids 1-24 and 359-428) are not resolved in the electron density map and thus are not modeled. The final atomic model contains amino acids 25-358 except for several disordered regions including an extracellular (amino acids 88-102) and two cytoplasmic segments (amino acids 161-195 and 318-323) (Fig. 2).

**Table 1.** Cryo-EM data collection, refinement and validation statistics.

|  | xPANX1<br>(EMD-xxxxxx)<br>(PDB xxxxx) | hPANX1<br>(EMD-xxxxxx)<br>(PDB xxxxx) |
| --- | --- | --- |
| **Data collection and processing** |  |  |
| Magnification | 105,000 | 105,000 |
| Voltage (kV) | 300 | 300 |
| Electron exposure (e-/Å<sup>2</sup>) | 62 | 62 |
| Defocus range (μm) | -1.0 to -2.5 | -1.0 to -2.5 |
| Pixel size (Å) | 1.1 | 1.1 |
| Symmetry imposed | C7 | C7 |
| Initial particle images (no.) | 765,110 | 745,589 |
| Final particle images (no.) | 176,371 | 15,796 |
| Map resolution (Å) | 3.38 | 3.77 |
| FSC threshold | 0.143 | 0.143 |
| Map resolution range (Å) | 3.0-5.0 | 3.0-6.0 |
| **Refinement** |  |  |
| Initial model used (PDB code) | 6NZZ | 6UZY |
| Model resolution (Å) | 3.40 | 3.70 |
| FSC threshold | 0.5 | 0.5 |
| Model resolution range (Å) | 3.20 | 3.50 |
| Map sharpening *B* factor (Å<sup>2</sup>) | -122 | -78.9 |
| Model composition |  |  |
| Non-hydrogen atoms | 16,212 | 11,739 |
| Protein residues | 1,946 | 1,463 |
| Ligands | 21 | 0 |
| *B* factors (Å<sup>2</sup>) |  |  |
| Protein | 47.8 | 64.6 |
| Ligand | 35.2 | N/A |
| R.m.s. deviations |  |  |
| Bond lengths (Å) | 0.008 | 0.005 |
| Bond angles (°) | 0.945 | 0.947 |
| Validation |  |  |
| MolProbity score | 1.53 | 1.33 |
| Clashscore | 6.98 | 6.03 |
| Poor rotamers (%) | 0 | 0 |
| Ramachandran plot |  |  |
| Favored (%) | 97.41 | 98.01 |
| Allowed (%) | 2.59 | 1.99 |
| Disallowed (%) | 0 | 0 |

**Figure 2.**
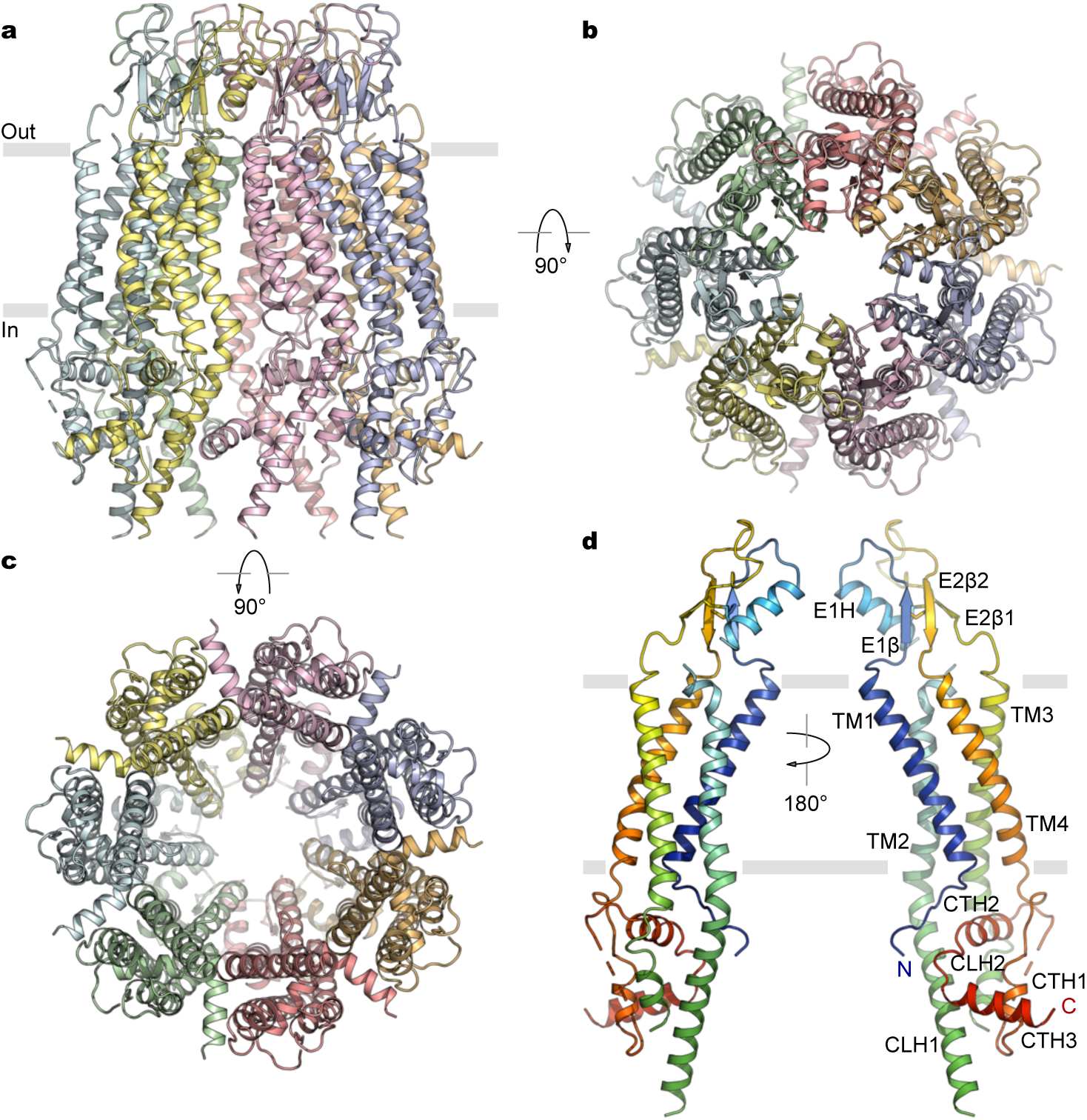
Structure of xPANX1. **a**-**c**, xPANX1 structure viewed parallel to the membrane (**a**), extracellularly (**b**), and intracellularly (**c**). Individual subunits are uniquely colored. Gray lines indicate the membrane boundary. **d**, Two views of a single subunit with its N- and C- termini, secondary structural elements and extracellular disulfide bonds indicated.

### Architecture of the heptameric PANX1 channel

The xPANX1 channel is assembled as a heptamer of seven identical subunits arranged around a central symmetry axis that constitutes the permeation pathway (Fig. 2a-c). Each subunit consists of a transmembrane domain with four membrane-spanning helices TM1-4 and folded extracellular and intracellular domains, which form the pore entrances. Both the N- and C-termini reside on the cytoplasmic side (Fig. 2d). Preceding the pore-lining helix TM1, an ordered N-terminal portion is located in the pore lumen and the distance between a pair of distal Cα atoms of the most N-terminal residues Y25 measures ∼35 Å. Following TM1, the first extracellular loop, containing a β-strand (E1β) and an α-helix (E1H), together with the second extracellular loop, creates a compact extracellular domain (ECD) structure, which comprises a three-stranded antiparallel β-sheet cross-linked to the E1H helix through a disulfide bond between C84 from E1H and C248 from E2β1 (Fig. 2d, Supplementary Fig. 4). The β-sheet is additionally strengthened by another disulfide bond between C66 from E1β and C267 from E2β2. The E1H helix contacts all three β-strands with its entirety and is sandwiched between two adjacent β-sheets, thus critically contributing to both intra- and inter-subunit interactions (Fig. 2a). On the intracellular side, two helices CLH1 and CLH2 from the cytoplasmic loop connecting TM2 and TM3 and three helices CTH1-3 from the C-terminal end following TM4 assemble into a compact cytoplasmic domain (CTD).

Seven ECDs are organized into a cap structure forming the extracellular entrance to the transmembrane pore, with the N-terminal end of each E1H helix protruding to the central pore axis (Fig. 2a, b). Below the extracellular cap, the pore-lining helix TM1 is tilted by ∼30° in the membrane and consequently the pore substantially widens toward the intracellular side, where TM2 begins to participate in lining the expanded pore (Fig. 2a, d). TM3 and TM4 are arranged at the periphery of the channel. The cytoplasmic helical domains extend away from the central pore axis, resulting in a voluminous intracellular vestibule (Fig. 2c). Following the C-terminal helix CTH3, the remaining C-terminal segment consisting of 70 amino acids, including the caspase cleavage sites^3,25^, is absent in the cryo-EM density map, suggesting that it is intrinsically dynamic. Thus, the structural and functional role of the C-terminus, which can be physiologically removed to promote channel activation^3,25^, remains undefined in the present structure. However, it is tempting to speculate that the distal C-terminal portion may be able to fold back and plug the pore as a means to facilitate channel closure, as suggested by previous studies^3,25,26^.

### Pore structure

Pore radius calculation reveals that the permeation pathway is exceptionally wide in the transmembrane and cytoplasmic portions and that the constriction site is generated by the ECDs (Fig. 3a, b). A ring of seven tryptophan residues (W74), each of which is located at the N-terminal end of the E1H helix, lines the wall of the outer pore with a diameter of ∼11 Å, potentially constituting an extracellular selectivity filter defining the maximum size of permeable molecules. The basic residue R75 from a neighboring E1H helix forms cation-π interaction with W74 and a salt bridge with D81 that is two helical turns away in the E1H helix, most likely stabilizing the tryptophan ring configuration. The network of inter-subunit electrostatic interactions, together with positive potential from the E1H helix dipole, results in an essentially neutral electrostatic potential at the extracellular entrance (Fig. 3c, d). Previous electrophysiology experiments have identified that W74 is the major determinant in CBX inhibition^33^, and that R75 is critical for inhibition by high concentrations of ATP and its analogues^34^. We introduced three alanine mutations at these positions (W74A, R75A, and D81A) and measured currents of mutant channels in excised membrane patches in the absence and presence of the inhibitor CBX (Supplementary Fig. 2c-e). D81A responds to CBX similarly as the wild type at negative membrane potentials but to a less extent at positive potentials. Remarkably, W74A is essentially insensitive to CBX and inhibition of R75A is much reduced at negative membrane potentials. These results, in addition to supporting our structural findings, reinforce the critical role of the tryptophan ring in channel pharmacology.

**Figure 3.**
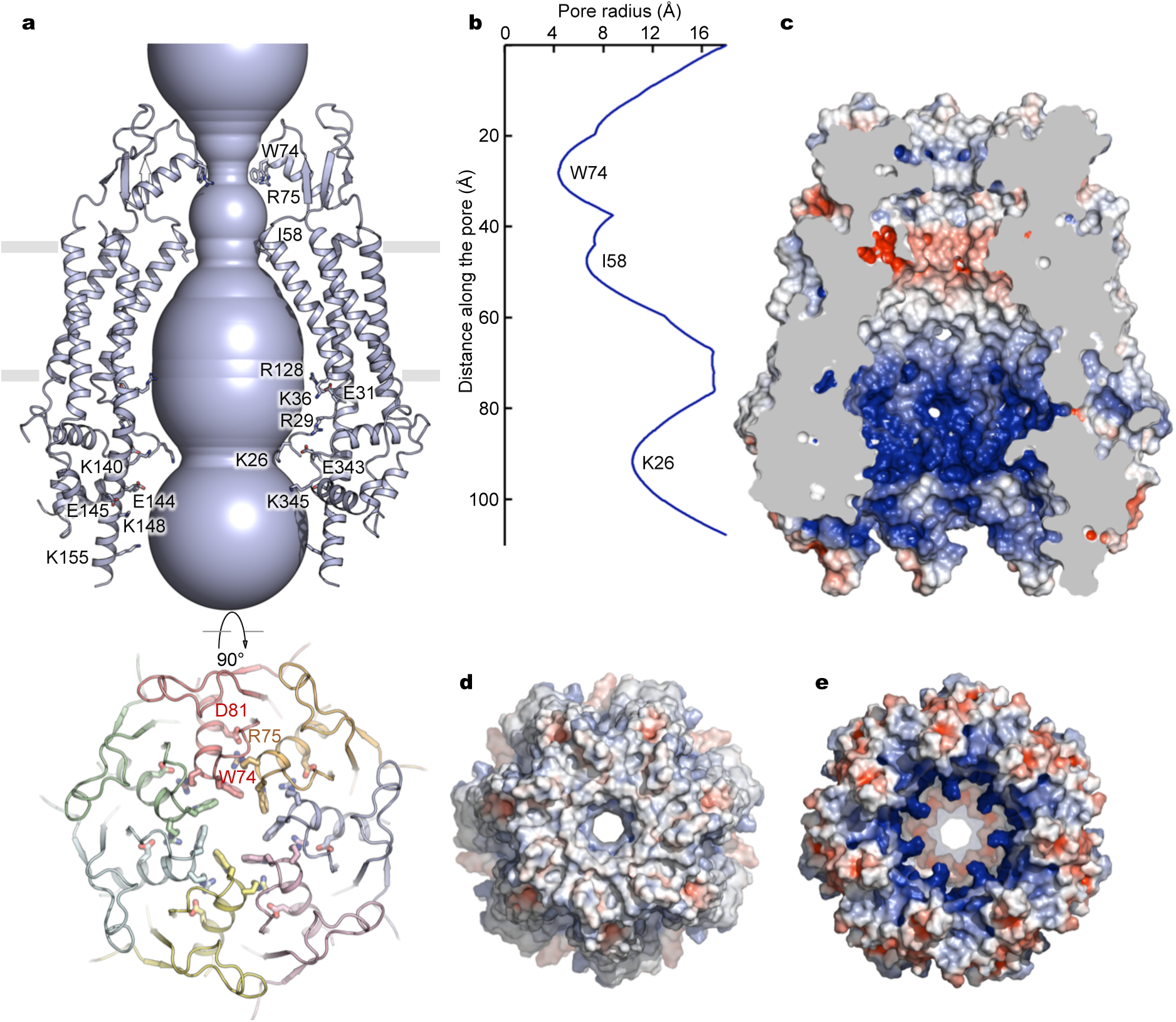
Pore architecture and electrostatic properties. **a**, The pore of xPANX1. Two opposing subunits are shown with selected residues lining the pore highlighted as sticks (top panel). View of the putative extracellular selectivity filter composed of a ring of seven tryptophan amino acids (W74) from the E1H helices (bottom panel). W74 forms a cation-π interaction with R75 from an adjacent subunit, which also forms a salt bridge with D81. **b**, The pore radius along the conduction path. **c**, Cutaway view of the pore lumen, colored by surface electrostatic potential (red, −5 kT/e; white, neutral; blue, +5 kT/e). **d, e**, Surface electrostatic potential of xPANX1 viewed from the extracellular (**d**) and intracellular (**e**) sides.

Below the extracellular entrance, the transmembrane pore is predominantly lined by hydrophobic amino acids from TM1 on the extracellular side (Fig. 3a, c). TM1 tilting in the membrane places its C-terminal extracellular end (I58) at the narrowest point of the pore in the transmembrane region, with a diameter of ∼17 Å. The N-terminal end of TM1 on the intracellular side is considerably distant from the central pore axis such that TM2 contributes to pore lining as well. The N-terminal tail preceding TM1 makes a turn and runs toward the central axis (Fig. 3a). Since the first N-terminal 24 amino acids are not resolved in the structure, it remains unknown whether the entire N-terminus stays in or departs from the pore lumen, and thus the pore dimensions in this region cannot be precisely determined. In the cytoplasmic vestibule, abundant charged amino acids face the pore lumen, generating an overall positive electrostatic potential, which in principle would facilitate recruitment of negatively charged ATP molecules to the intracellular entryway to the transmembrane pore (Fig. 3c, e).

The PANX1 structure shows resemblance to structures of the extended 4-TM channel family members including intercellular gap junction channels connexins and innexins, and the more recently identified volume-regulated anion channels formed by leucine-rich repeat-containing protein 8 (LRRC8) family members^35–37^, despite low amino acid sequence similarity between these channel proteins (Fig. 4a-f). In PANX1 and LRRC8, the E1H helix in the first extracellular loop creates the narrowest constriction at the outer pore region, whereas in connexin and innexin hemichannels, the N-terminal helix defines the narrowest constriction within the transmembrane portion of the pore (Fig. 4b-f). Interestingly, analogous to W74 in PANX1 forming the extracellular constriction, a corresponding arginine residue (R103) at the N-terminal end of E1H in LRRC8A generates a narrower extracellular constriction, likely functioning as an anion selectivity filter^37^ (Fig. 4b-d). Despite the constriction sites being differently organized, the overall pore dimensions of PANX1 are comparable to those of the open connexin hemichannel (Fig. 4b, c, e), suggesting that our PANX1 structure represents an open conformation and that PANX1 and connexin may have similar size restrictions for permeable substrates. Interestingly, these channels are assembled from different numbers of subunits, though each channel subunit adopts a similar structural fold. While the connexin hemichannel and LRRC8 are hexameric, PANX1 is heptameric and the innexin hemichannel is octameric.

**Figure 4.**
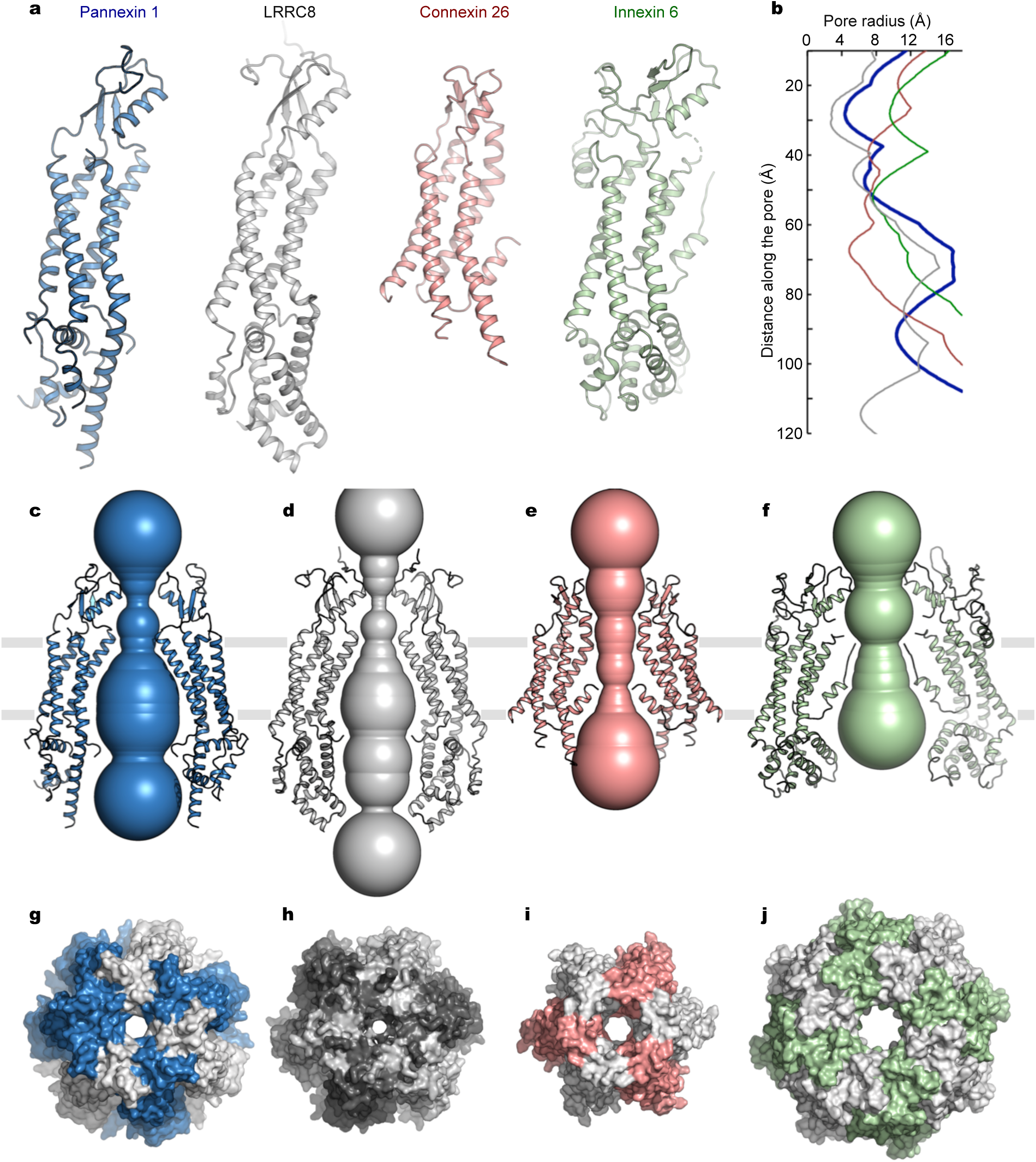
Structural comparison of the extended 4-TM family of channels. **a**, Subunit structures of the pannexin 1 (blue), LRRC8 (gray, PDB: 6G8Z), connexin 26 (red, PDB: 2ZW3), and innexin 6 (green, PDB: 5H1Q) channels. **b**, Pore radii of the pannexin 1 (blue), LRRC8 (gray), connexin 26 (red), and innexin 6 (green) channels. **c**-**f**, Ion-conduction pores of the pannexin 1 (**c**), LRRC8 (**d**), connexin 26 (**e**), and innexin 6 (**f**) channels. Only two opposing subunits are shown for each channel. **g**-**j**, Surface representation of the heptameric pannexin 1 (**g**), hexameric LRRC8 (**h**), hexameric connexin 26 (**i**), and octameric innexin 6 (**j**) channels. Channel subunits are in alternate colors.

To substantiate that PANX1 channels are indeed heptameric and that the heptameric assembly is not singularly specific for the frog ortholog, we extended our structural analysis to human PANX1 (hPANX1). We purified the full-length wild-type hPANX1 channel and conducted cryo-EM analysis. Reference-free 2D classification clearly confirmed the heptameric channel assembly and 3D reconstruction yielded a structure at a lower resolution of ∼3.8 Å (Fig. 5, Supplementary Fig. 5). Notably, the cytoplasmic domain in the hPANX1 structure is largely disordered. Nonetheless, the overall structure and dimensions of the extracellular constriction and the transmembrane pore of hPANX1 and xPANX1 are similar (Fig. 5). Thus, we have ascertained that PANX1 channels are heptameric through structural characterization of two orthologs, and in this article focus on the higher-resolution, more complete atomic model of xPANX1.

**Figure 5.**
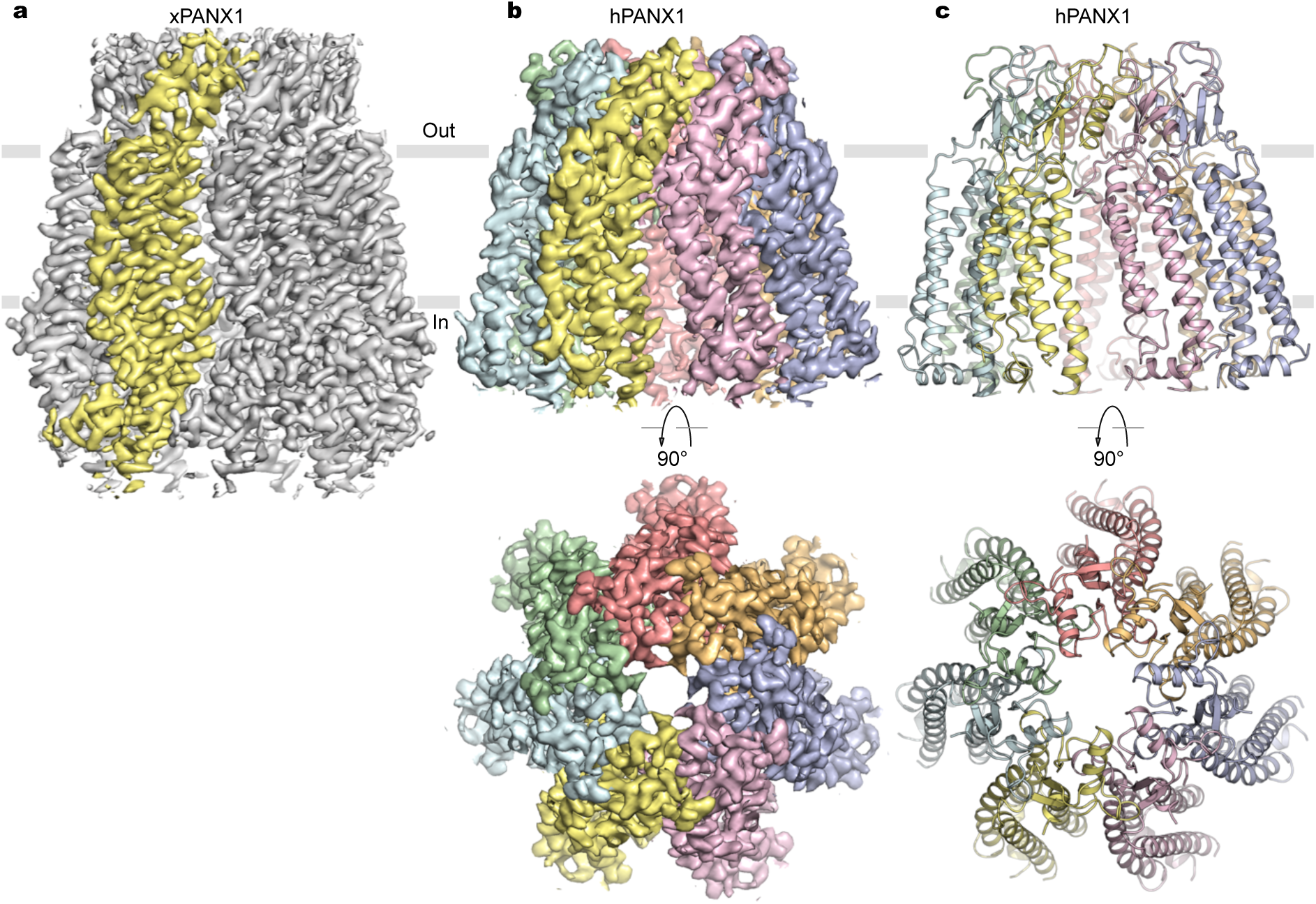
Comparison of cryo-EM reconstructions of human and frog PANX1. **a**, Cryo-EM density map of xPANX1 contoured at 6.0 σ. One of the subunits is colored in yellow. **b**, Orthogonal views of cryo-EM density map of human PANX1 contoured at 6.0 σ. Each subunit is in a unique color. **c**, Orthogonal views of the heptameric structure of human PANX1.

### Inter-subunit interface

The inter-subunit interface in the heptameric xPANX1 channel, contributed by the extracellular, transmembrane, and intracellular components, buries ∼3,900 Å^2^ of molecular surface per subunit. The interface in the transmembrane portion is primarily mediated by hydrophobic interactions between TM2 and TM1. Pronounced non-polypeptide electron densities attributable to membrane lipids based on the shape, location and chemical environment fill in the crevice between TM3 and an adjacent TM4 at the periphery of the channel (Fig. 6a). In particular, head-and-two-tails shaped density is observed at the inner leaflet of the lipid membrane where channels naturally reside, and the putative lipid head group is surrounded by polar amino acids including positively charged residues from the N-terminal end of TM3 (K214) and the C-terminal domain immediately following TM4 (R302, K303) (Fig. 6b). It appears that the lipid acyl chains enhance inter-subunit packing within the transmembrane domains, while the lipid head groups interact with polar and charged residues in the cytoplasmic domains. Thus, these lipids essentially function as molecular glues to augment interactions between subunits as well as coupling between the transmembrane and cytoplasmic domains (Fig. 6b).

**Figure 6.**
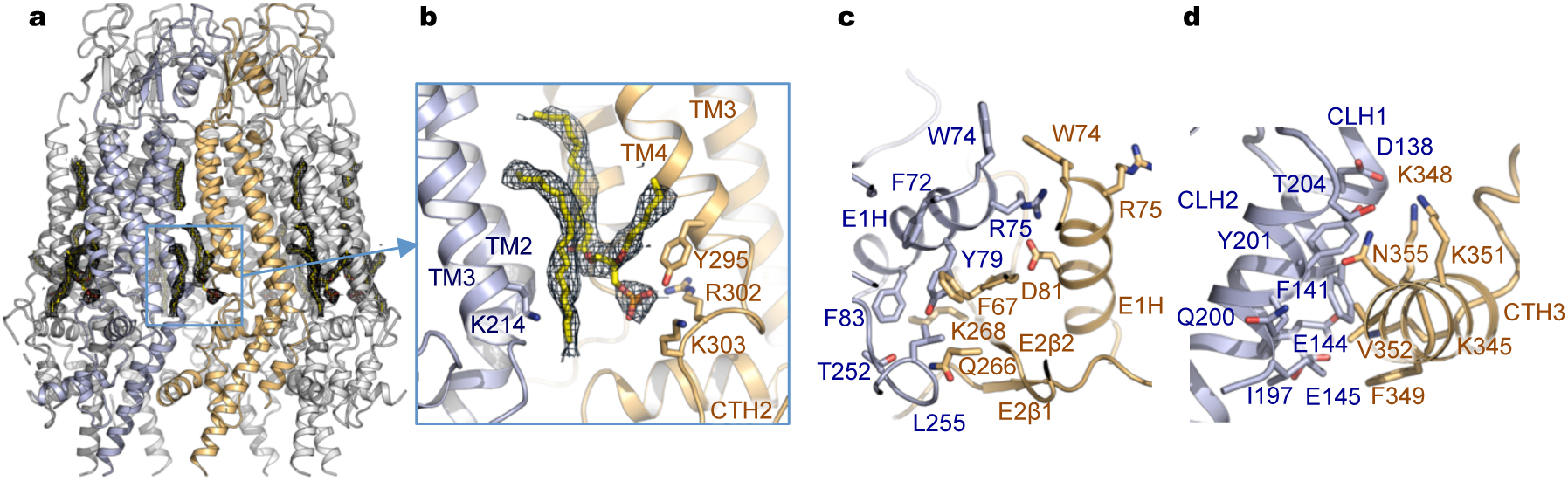
Inter-subunit interface. **a**, Robust lipid-like densities at the outer and inner leaflets located at the crevice between channel subunits. The lipid electron density is contoured at 4.0 σ. **b**, Detailed view of the inner leaflet lipids. A putative phospholipid is modeled to illustrate potential interactions with surrounding polar residues from the channel. **c**,**d**, The extracellular (**c**) and intracellular (**d**) inter-subunit interfaces. Selected residues involved in salt bridge, hydrogen bonding or hydrophobic interactions are highlighted, with side chains shown in stick representation.

On the extracellular side, the W74-R75-D81 inter-subunit interaction network from the E1H helices contributes to the outer pore constriction. In addition, multiple contact points from other structural elements of the ECDs are involved in the inter-subunit interface, such as F67-Y79 aromatic-aromatic and Q266-T252 hydrogen bonding (Fig. 6c). These specific side chain interactions between the ECDs probably further support the outer pore assembly. On the intracellular side, the last resolved C-terminal helix CTH3 engages an extensive network of interactions with the cytoplasmic linker helices CLH1 and CLH2 from an adjacent subunit (Fig. 6d). In line with this strategic arrangement, two human mutations (K346E and C347S) from CTH3 have been reported to increase channel activity, resulting in female infertility associated with oocyte death^15^. In particular, the corresponding residue K348 in xPANX1, together with K351, forms salt-bridge interactions with D138 from CLH1. Charge reversal by the human K346E mutation would disturb this interface. These structural and functional observations suggest that the cytoplasmic interface plays an important role in controlling channel activity.

## Discussion

Despite the increasing recognition of physiological and pathophysiological functions of PANX1 channels as well as an accumulating body of literature, understanding of channel properties remains rather primitive and consensus is still lacking regarding fundamental characteristics, such as gating, permeability, single-channel conductance, and number of activated states, perhaps owing to the intrinsic complexity of the channel under different cellular contexts^1,2^. High-resolution 3D structures representing distinct functional states are essential for resolving discrepancies and elucidating the inner workings of the channel. Toward this end, we have determined cryo-EM structures of PANX1 at near atomic resolution, delineating the general architecture and atomic detail of an important family of ATP-release channels that were previously unknown. Our human and frog PANX1 structures now unambiguously establish that PANX1 channels are heptameric. Supporting its role in ATP permeation, PANX1 has an ample central conduction pore with dimensions comparable to those of intercellular gap junction connexin channels, which conduct ions as well as large molecules such as ATP. The open PANX1 structures, in conjunction with a small unitary conductance in excised membrane patches, suggests that ATP release does not necessarily require a large unitary conductance.

In the absence of a closed conformation, locations of the channel gates and structural rearrangements associated with channel closure remain to be elucidated. Scanning cysteine accessibility experiments and mutagenesis data have suggested that the distal C-terminal tail resides within the pore to inhibit naive channels^25,38^, that dissociation of the C-terminal tail immediately following the caspase cleavage site from the channel is necessary for activation^25^, and that sequential removal of the C-terminal tails in the oligomeric channel results in stepwise activation with graded increase in single-channel conductance and permeant size^26^. Consistent with these notions, the inhibitory C-terminal tail, potentially functioning as a mobile pore blocker^25,26^, is not observed in the pore lumen of the open PANX1 structures. However, channels with the C-terminal tails fully removed still transition to closed states^26^, suggesting additional gates at other locations of the channel.

The putative selectivity filter formed by the conserved tryptophan ring (W74) from the E1H helices in the first extracellular loops could potentially operate as a gate. CBX inhibits PANX1 primarily through interactions with W74 and nearby residues in the first extracellular loop, as demonstrated in our electrophysiology experiments as well as previous studies^33^. These results suggest that the selectivity filter and its immediate surrounding could indeed function as a gate. The N-terminal tail preceding TM1 resides in the intracellular pore lumen and the first 24 amino acids are not resolved in the open structure. Conceptually the N-terminal tail could extend to the central pore axis, constituting an intracellular gate. These possibilities need to be addressed with additional structural studies capturing distinct functional states of the channel.

## Methods

### Cloning, expression and purification of PANX1 channels

DNA fragments encoding *Homo sapiens* (hPANX1, NCBI: NP_056183.2) and *Xenopus tropicalis* pannexin 1 (xPANX1, NCBI: NP_001123728.1) were synthesized (Gene Universal) and cloned into a modified yeast *Pichia pastoris* expression vector pPICZ-B with a C-terminal GFP-His_10_ tag linked by a PreScission protease cleavage site. For electrophysiological recordings, the corresponding DNA fragments were ligated into a modified pCEU vector containing a C-terminal GFP-His_8_ tag. Mutations were introduced by site-directed mutagenesis.

For protein purification, yeast cells were disrupted by milling (Retsch MM400) and resuspended in lysis buffer containing 50 mM Tris pH 8.0 and 150 mM NaCl supplemented with protease inhibitors (2.5 µg/ml leupeptin, 1 µg/ml pepstatin A, 100 µg/ml 4-(2-Aminoethyl) benzenesulfonyl fluoride hydrochloride, 3 µg/ml aprotinin, 1 mM benzamidine and 200 µM phenylmethane sulphonylfluoride) and DNase I. Cell membranes were solubilized in buffer containing 1% (w/v) Lauryl Maltose Neopentyl Glycol (LMNG, Anatrace) for 2 h with stirring at 4°C. Solubilized PANX1 protein was separated from the insoluble fraction by centrifugation for 1 h at 30,000 g, and incubated with 3 ml of cobalt-charged resin (G-Biosciences) for 3 h at 4°C. Resin was then washed with 10 column volumes of buffer containing 20 mM Tris pH 8.0, 150 mM NaCl, 10 mM imidazole, and 85 µM glyco-diosgenin (GDN, Anatrace). The C-terminal GFP-His_10_ tag was removed by digestion with PreScission protease at 4°C overnight on the column. The flow-through was then collected and concentrated using a 100 kDa concentrator and further purified on a Superose 6 gel filtration column (GE Healthcare Life Sciences) in 20 mM Tris pH 8.0, 150 mM NaCl and 40 µM GDN. The fractions corresponding to the oligomeric channels were collected and concentrated to ∼6 mg/ml for cryo-EM grid preparation.

### Cryo-EM sample preparation and imaging

Cryo-EM grids were prepared using FEI Vitrobot Mark IV (FEI). 3.5 µl of purified channel proteins at ∼6 mg/ml were pipetted onto glow-discharged copper Quantifoil R2/2 holey carbon grids (Quantifoil). Grids were blotted for 2 s at ∼100% humidity and flash frozen in liquid ethane. The grids were loaded into a Titan Krios (FEI) electron microscope operating at 300 kV with a Gatan K2 Summit (Gatan) detector. GIF Quantum energy filter with a slit width of 20 eV was operated in zero-energy-loss mode before detector. Images were recorded with the EPU software (https://www.fei.com/software/epu-automated-single-particles-software-for-life-sciences/) in the counting mode with a pixel size of 1.1 Å and a nominal defocus value ranging from −1.0 to −2.5 µm. Data were collected with a dose of ∼7.8 electrons per Å^2^ per second, and each movie was recorded with an 8 s exposure and 40 total frames (200 ms per frame), resulting in an accumulated dose of ∼62 electrons per Å^2^.

### Image processing and map calculation

Recorded movies were first aligned and dose-weighted with MotionCor2^39^, and then subjected to contrast transfer function (CTF) determination using GCTF^40^. Following motion correction and CTF estimation, low-quality images were manually removed from the datasets. For the xPANX1 dataset, 1903 particles were manually selected to generate two-dimensional class templates for automated particle picking in RELION3^41^. Template-based automatic picking resulted in 765,110 particles from 1,823 micrographs. Particles were extracted using a box size of 240 pixels and subjected to 2D classification with a mask diameter of 180 Å. Low-quality particle images were discarded by performing two rounds of 2D classification. The resulting good 2D classes containing 272,279 particles were selected and imported into cryoSPARC^42^ to generate an initial map for 3D classification in RELION3^41^. After 3D classification, two classes showing intact channel features (176,371 particles) were selected and subjected to 3D refinement and post-processing, yielding an overall resolution of 3.43 Å. CTF refinement and Bayesian polishing were performed in RELION3 to further improve the resolution to 3.38 Å.

Data processing for hPANX1 followed a similar procedure. Briefly, 7,469 particles were picked using LoG-based auto-picking to generate 2D classes in RELION3. After several rounds of 2D classification, 2D classes with a total of 5,048 particles representing different orientations of the channel were selected as templates for automated particle picking. A total of 745,589 particles from auto-picking were subjected to two rounds of 2D classification, yielding 275,151 good particles. 3D classification was carried out using a low-pass-filtered xPANX1 map with C7 symmetry imposed. One of the 3D classes with 55,848 particles showing channel features was subjected to 3D refinement, reaching a nominal resolution of 4.55 Å. 3D classification without particle alignment was then performed. Further 3D refinement and post-processing of one of the classes (15,796 particles) resulted in an overall resolution of 3.88 Å. Three rounds of CTF refinement and Bayesian polishing were subsequently performed to further improve the resolution to 3.77 Å. Local resolution estimates were calculated in RELION3^41^.

### Model building and coordinate refinement

A homology model of xPANX1 was generated using coordinates of mouse LRRC8A (PDB: 6NZZ) by the SWISS-MODEL server^43^. The model was placed into the cryo-EM density map using UCSF Chimera^44^. The TM1-4, E1H, CLH1 helices as well as the β sheet containing E1β, E2β1 and E2β2 were further adjusted to fit the density, and the remaining model was *de novo* built in COOT^45^. Bulky side chains and disulfide bonds were used to facilitate sequence register. Cycles of model building in COOT and refinement using real_space_refine in PHENIX^46^ were performed to obtain the final refined model. For hPANX1, a homology model was first generated by the SWISS-MODEL server using the refined model of xPANX1, and then fitted into the cryo-EM density map in UCSF Chimera. Following rigid body refinement in PHENIX, cycles of model building in COOT and refinement using real_space_refine in PHENIX were performed. The final refined models were validated using MolProbity^47^. Pore dimensions were calculated using the program HOLE^48^. Structural figures were made using USCF Chimera^44^ and PyMol (pymol.org).

### Electrophysiological recordings

CosM6 cells were transfected with ∼1 µg of wild type or mutant constructs using FuGENE6 (Promega) and were patched within 1-2 days after transfection. Symmetric internal sodium (150 mM NaCl, 1 mM EGTA, 1 mM EDTA, 10 mM Hepes, pH 7.4) or potassium (150 mM KCl, 1 mM EGTA, 1 mM EDTA, 10 mM Hepes, pH 7.4) buffers were used for all recordings. Upon patch excision in the inside-out configuration, the bath was continuously perfused with symmetric buffer with or without 100 µM carbenoxolone disodium salt (CBX, Alfa Aesar), so that the drug was always applied only from the cytoplasmic side of the membrane. Recordings were made and digitized with the Axopatch 1D patch-clamp amplifier and the Digidata 1320 digitizer (Molecular Devices) and analyzed with the pClamp software suite (Molecular Devices). The pipettes with bubble number (BN)^49^ 3.5-5.0 were made from the Kimble Chase 2502 soda lime glass using a Sutter P-86 puller (Sutter Instruments).

For single channel recordings, the data were collected at 15 kHz and low-pass filtered at 5 kHz. The measurements were carried out at −60 mV membrane potential. The pipettes with BN ∼3.5-4.0 were used. For spontaneous activation recordings, the data were collected at 500 Hz and low-pass filtered at 200 Hz. The measurements were carried out at −30 mV membrane potential. The pipettes with BN ∼5.0 were used. For current-voltage (IV) relationship recordings, the data were collected at 1 kHz and low-pass filtered at 500 Hz. The measurements were carried out using 2 s voltage ramps from 100 mV to −100 mV. The measurement protocol performed repetitive recordings of the response to a voltage ramp every 10 s until current saturation, at which point CBX was added. The pipettes with BN ∼5.0 were used.

## Data availability

The cryo-EM maps of hPANX1 and xPANX1 have been deposited to Electron Microscopy Data Bank with accession code EMD-xxxxx and EMD-xxxxx. Atomic coordinates for hPANX1 and xPANX1 structures have been deposited to the Protein Data Bank (PDB) with accession code xxxx and xxxx. Other source data are available from the corresponding authors upon request.

## Acknowledgements

This work was supported by start-up funds from Washington University School of Medicine (to P.Y.). M.J.R and J.A.J.F are supported by the Washington University Center for Cellular Imaging, which is funded, in part by Washington University School of Medicine through the Precision Medicine Initiative, the Children’s Discovery Institute of Washington University and St. Louis Children’s Hospital (CDI-CORE-2015-505 and CDI-CORE-2019-813) and the Foundation for Barnes-Jewish Hospital (3770).

## Author Contributions

Z.D., Z.H and R.M.B. performed biochemical preparations, cryo-EM experiments, structural determination and analysis. G.M. conducted electrophysiology experiments. M.R., J.A.J.F., Z.D. and Z.H. performed cryo-EM data acquisition. P.Y. designed and supervised the project. Z.D., Z.H., G.M., and P.Y. analyzed the results and prepared the manuscript with input from all authors. Correspondence and requests for materials should be addressed to P.Y.

---

**Supplementary Figure 1.**
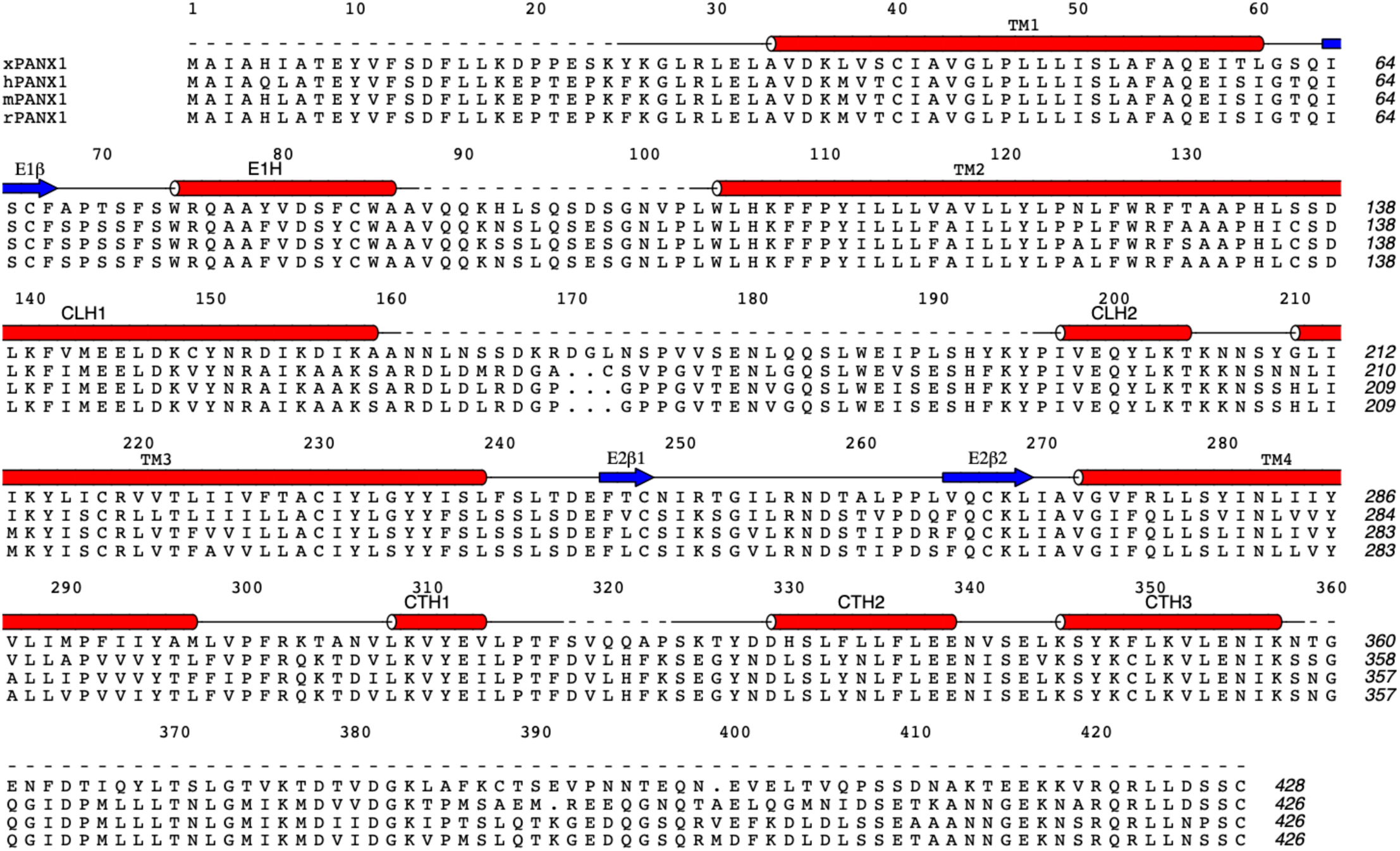
Sequence alignment of PANX1 orthologs. The protein sequences of *Xenopus tropicalis* (xPANX1, NCBI sequence: NP_001123728.1), *Homo sapiens* (hPANX1, NCBI sequence: NP_056183.2), *Mus musculus* (mPANX1, NCBI sequence: NP_062355.2), and *Rattus norvegicus* PANX1 (rPANX1, NCBI sequence: NP_955429.1) are aligned and secondary structural elements of xPANX1 are shown above the protein sequences. Dashed lines indicate unresolved regions in the xPANX1 structure.

**Supplementary Figure 2.**
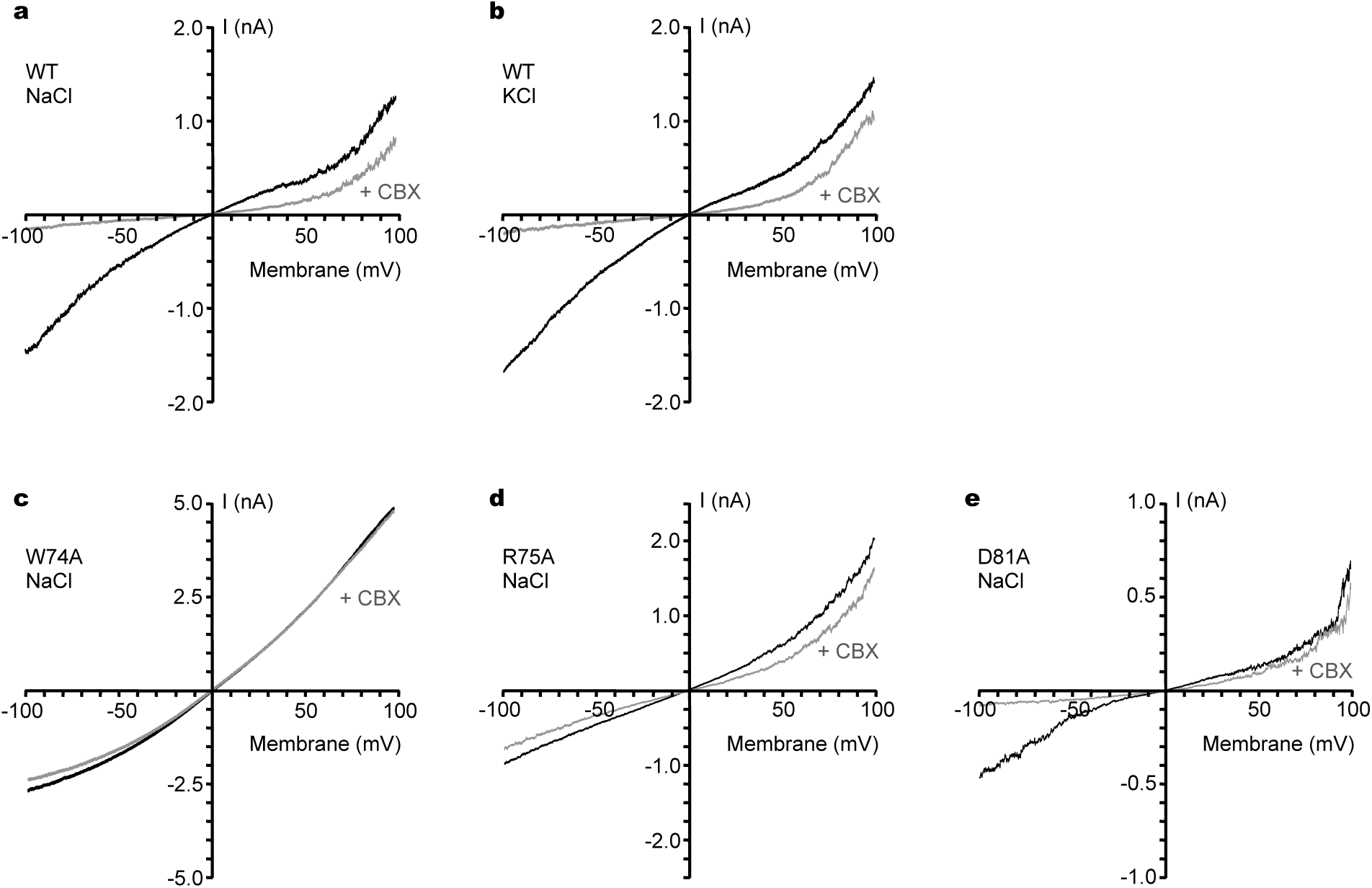
CBX inhibition of the wild-type xPANX1 and mutants in excised inside-out membrane patches. **a**,**b**, Current-voltage relationship of wild-type xPANX1 in symmetrical NaCl (**a**) and KCl (**b**) in the absence (black) and presence (gray) of 100 µM CBX. **c**-**e**, Current-voltage relationship of the W74A (**c**), R75A (**d**), and D81A (**e**) mutants in symmetrical NaCl in the absence (black) and presence (gray) of 100 µM CBX.

**Supplementary Figure 3.**
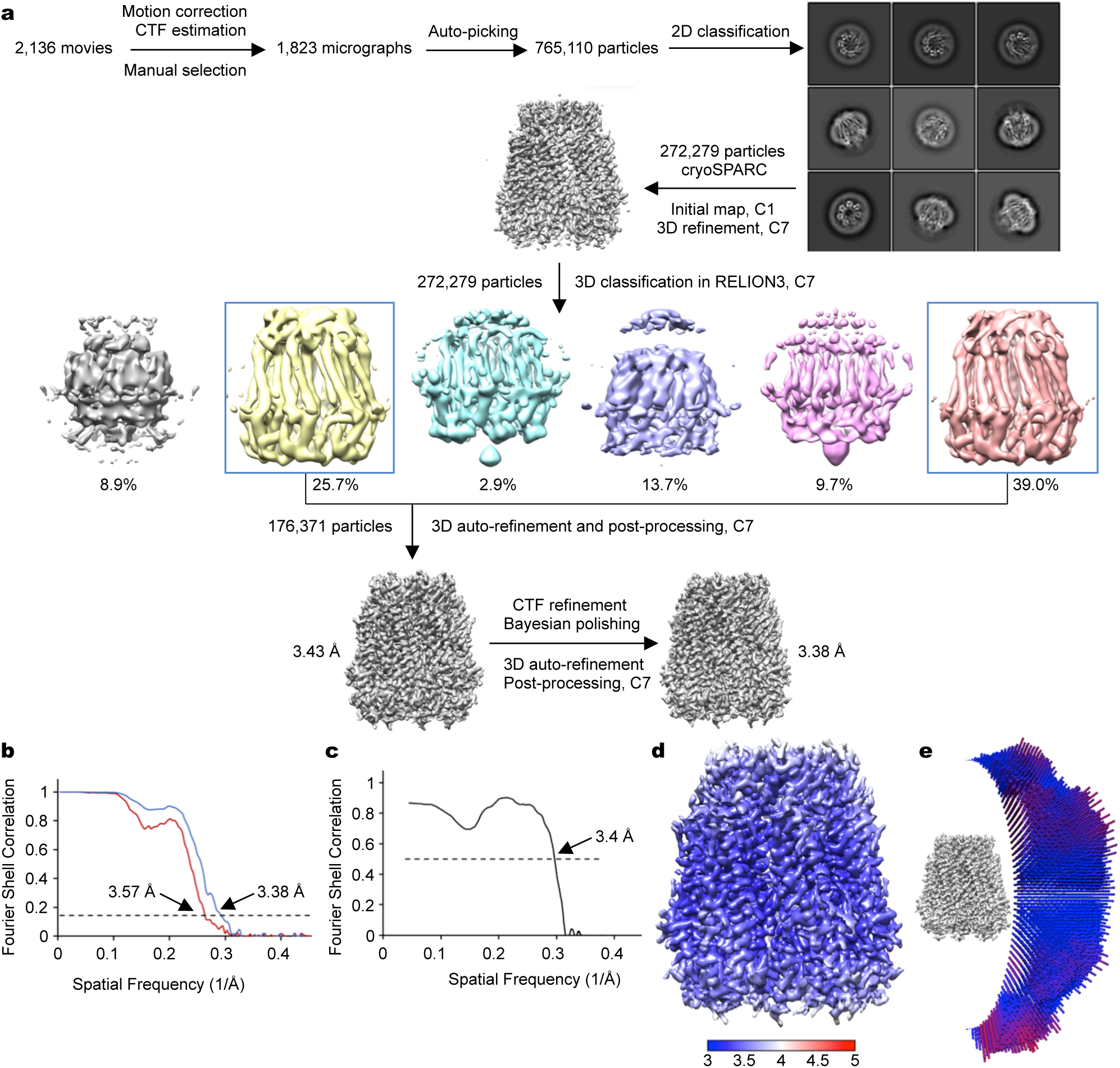
Cryo-EM reconstruction of xPANX1. **a**, Flowchart of cryo-EM image processing. **b**, Fourier shell correlation before and after post-processing in RELION3. **c**, Fourier shell correlation between the refined model and the full map. **d**, Cryo-EM density map colored by local resolution in the range of 3.0-5.0 Å. **e**, Angular distribution plot of particles used in the final reconstruction. Only one-seventh of the sphere is shown owing to applied C7 symmetry.

**Supplementary Figure 4.**
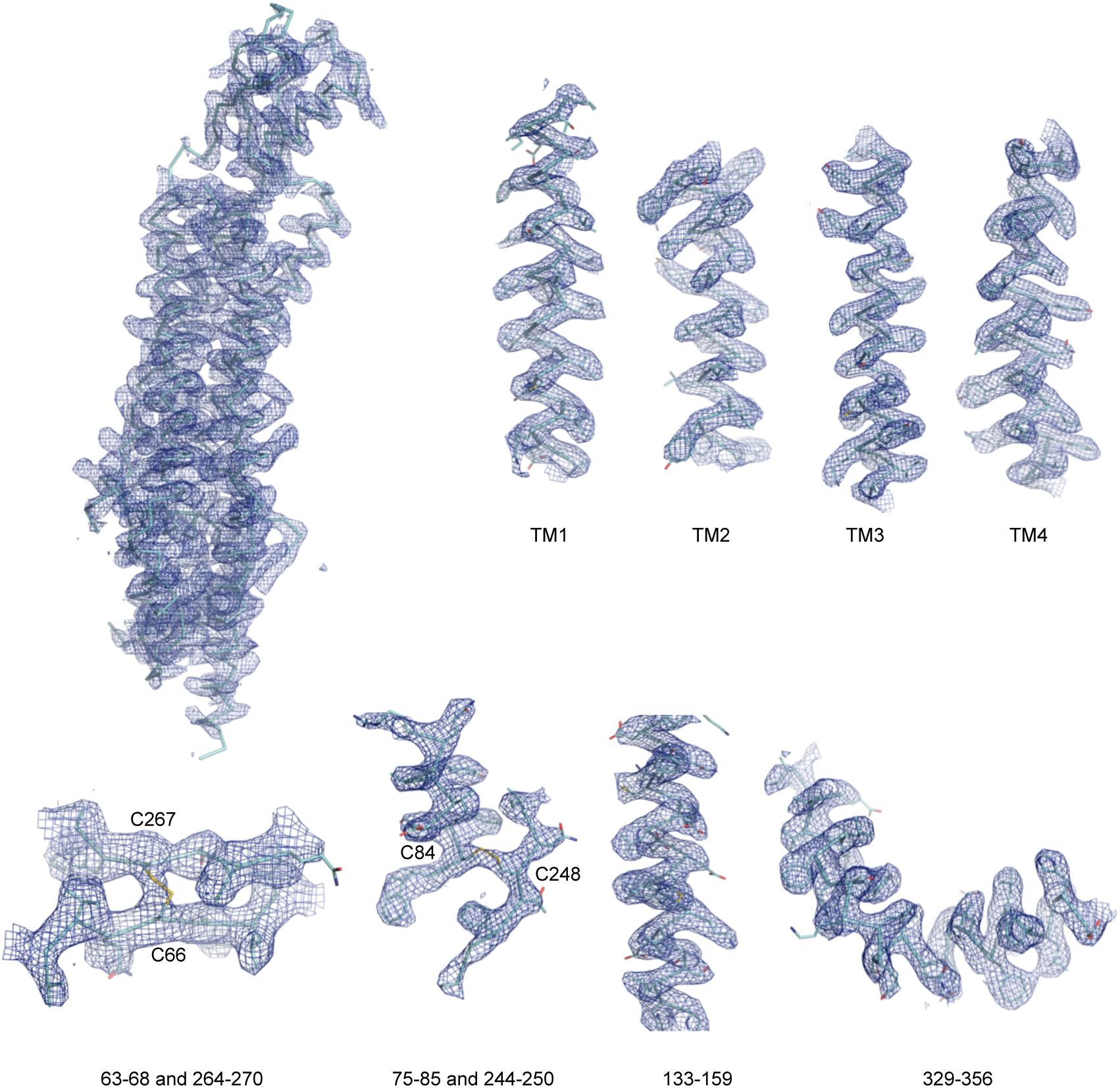
Cryo-EM density map of xPANX1. Electron density for an entire channel subunit as well as for selected regions is shown as blue mesh. Residues forming disulfide bonds are highlighted.

**Supplementary Figure 5.**
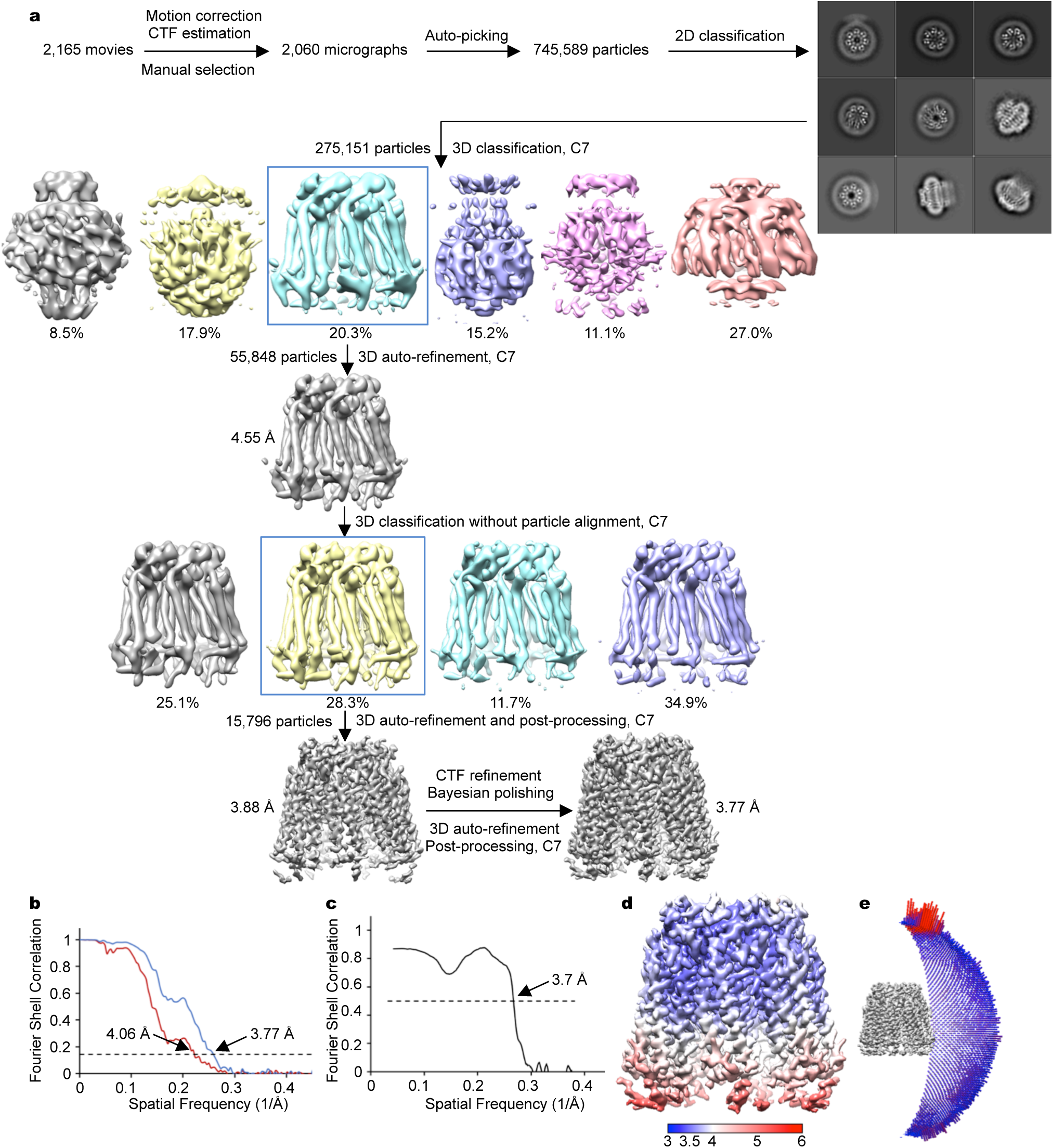
Cryo-EM reconstruction of human PANX1. **a**, Flowchart of cryo-EM image processing. **b**, Fourier shell correlation before and after post-processing in RELION3. **c**, Fourier shell correlation between the refined model and the full map. **d**, Cryo-EM density map colored by local resolution in the range of 3.0-6.0 Å. **e**, Angular distribution plot of particles used in the final reconstruction.

